# Distinct effects of nonselective Rho-kinase inhibitor fasudil and selective Rho-kinase 2 inhibitor KD025 on serotonin and dopamine release in the nucleus accumbens of mice

**DOI:** 10.1101/2025.07.21.666024

**Authors:** Rinako Tanaka, Taku Nagai, Toshitaka Nabeshima, Kozo Kaibuchi, Norio Ozaki, Hiroaki Ikesue, Hiroyuki Mizoguchi, Kiyofumi Yamada

## Abstract

Recent studies have indicated that the Rho GTPase family and Rho-kinases are associated with psychiatric diseases, such as schizophrenia. Rho-kinases have two subtypes, Rho-kinases 1 and 2 that regulate actin dynamics and mediate neurite outgrowth, spine morphology in neurons, and neurotransmitter release in vitro and ex vivo. However, the precise role of Rho-kinases in neurotransmitter release in vivo remains unclear. To clarify the role of Rho-kinases 1 and 2 in serotonin and dopamine release in the nucleus accumbens (NAc) of mice in vivo, we investigated the effect of a nonselective Rho-kinase inhibitor, fasudil, and a selective Rho-kinase 2 inhibitor, KD025, using an in vivo microdialysis technique. Fasudil perfusion (1–20 μM) into the NAc increased the basal extracellular serotonin level but did not affect dopamine levels, whereas KD025 (10–20 μM) had little effect on basal serotonin and dopamine levels. Notably, fasudil perfusion into the NAc suppressed depolarization-induced serotonin and dopamine release in a dose-dependent manner, whereas KD025 selectively suppressed depolarization-induced serotonin release. Our results suggested that Rho-kinases 1 and 2 are associated with dopamine and serotonin release, respectively, and that both may have significant but distinct roles in the regulation of serotonin and dopamine release in the NAc.

## Introduction

Neurotransmitters, such as serotonin and dopamine, are crucial for normal brain functioning but are also associated with the pathophysiology of mental diseases, such as schizophrenia, depression, and autism spectrum disorders [1–5]. Recently, the Rho GTPase family and Rho-kinases have been associated with schizophrenia, depression, autism spectrum disorders, and intellectual disabilities [6–12]. Rho-kinase is a serine/threonine kinase that plays a role in regulating actin dynamics and mediating neurite outgrowth and neuronal spine morphology [13–16].

Many studies have reported that Rho-kinase is involved in the regulation of neurotransmitter release in vitro and ex vivo [15,17–24]. However, the results of these studies were controversial, and the role of Rho-kinase in the regulation of neurotransmitter release in vivo remains undetermined.

Rho-kinases have two subtypes, Rho-kinases 1 and 2 [14,25], which show similar kinase activities but different distributions in the brain [25–27]. Indeed, knockout of these subtypes results in different spine morphology profiles in mice [28]. Thus, Rho-kinases 1 and 2 may play distinct roles in neurotransmitter release. In the present study, to clarify the roles of Rho-kinases 1 and 2 in neurotransmitter release in vivo, we investigated whether the nonselective Rho-kinase inhibitor fasudil and the selective Rho-kinase 2 inhibitor KD025 modulate serotonin and dopamine release in the nucleus accumbens (NAc) using an in vivo microdialysis technique.

## Materials and methods

### Animals

Male mice were used to exclude any potential estrous cycle effects in female mice [29]. Male C57BL/6J mice (n = 46) were obtained from the Japan SLC (Shizuoka, Japan). Male mice aged 8–9 weeks were used in this study. The mice were housed in groups of four to six mice per cage (28 cm length × 17 cm width × 13 cm height) in standard conditions (23 ± 1 °C, 50 ± 5% humidity) with a 12-h light/dark cycle (9:00 am – 9:00 p.m.). Food and water were provided ad libitum. Animals were handled in accordance with the guidelines established by the Institutional Animal Care and Use Committee of Nagoya University, Guiding Principles for the Care and Use of Laboratory Animals approved by the Japanese Pharmacological Society, and Guide for the Care and Use of Laboratory Animals by the National Institutes of Health of the United States.

### Drug treatment

Fasudil monohydrochloride salt (99% purity) was supplied by Asahi Kasei Pharma Corp. (Tokyo, Japan). Fasudil and KD025 (Slx-2119) (Cat#HY-15307/CS-0776; MedChemExpress, Monmouth Junction, NJ, USA) were suspended in distilled H_2_O and dimethyl sulfoxide, respectively, as stock solutions. Stock solutions were suspended in artificial cerebrospinal fluid (aCSF; 147 mM NaCl, 4 mM KCl, and 2.3 mM CaCl_2_). The fasudil and KD25 aCSF controls contained distilled H_2_O (0.3%) and dimethyl sulfoxide (0.2%), respectively.

### In vivo microdialysis

Mice were anesthetized using medetomidine (0.3 mg/kg), midazolam (4 mg/kg), and butorphanol (5 mg/kg), and a guide cannula (AG-6, Eicom Corp., Kyoto, Japan) was implanted in the NAc (+1.5 mm anteroposterior, +0.8 mm mediolateral from the bregma, and 4.0 mm dorsoventral from the skull) according to the mouse brain atlas [30] (Fig. 1). Upon recovery from surgery, a dialysis probe (FX-I-6-01; membrane length of 1 mm, Eicom Corp.) was inserted through the guide cannula and perfused with aCSF at a flow rate of 1.0 µL/min. The outflow fractions were collected every 5 min. After collecting the baseline fractions, mice were treated with fasudil (1, 10, or 20 μM), KD025 (10 or 20 μM), or a vehicle-containing aCSF through the dialysis probe, and drug treatment was continued until the end of the experiment (Figs. 1 and 2). To examine the effects of Rho-kinase inhibitors on depolarization-induced serotonin and dopamine release, high-K^+^-containing aCSF (60 mM; isomolar replacement of NaCl with KCl) was perfused through the dialysis probe for 20 min, and the aCSF was changed to 4 mM KCl-containing normal aCSF (Figs. 3 and 4). Serotonin and dopamine levels in the dialysates were analyzed using a high-performance liquid chromatography system (HTEC-500, Eicom Corp.) equipped with an electrochemical detector.

**Figure 1.**
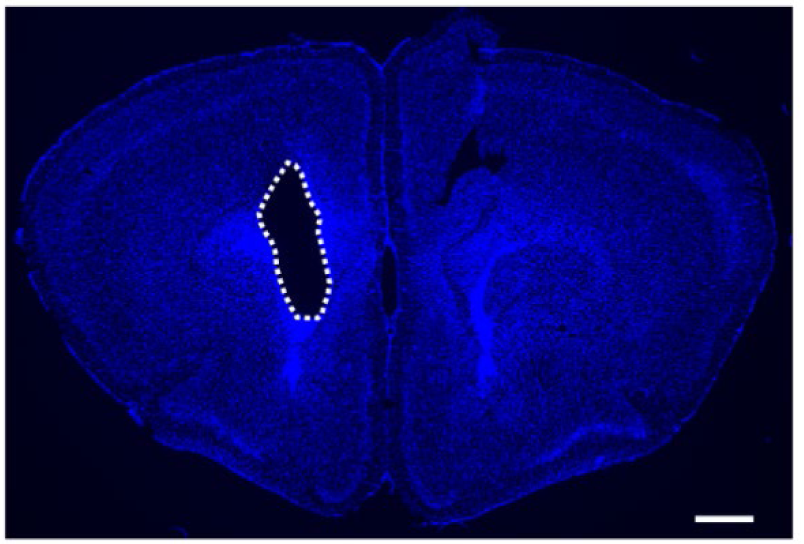
Representative image of the guide cannula trace in the NAc Blue signals represent Hoechst 33342. White dashed line represents trace of guide cannula (scale bar indicates 500 μm). NAc, Nucleus accumbens

**Figure 2.**
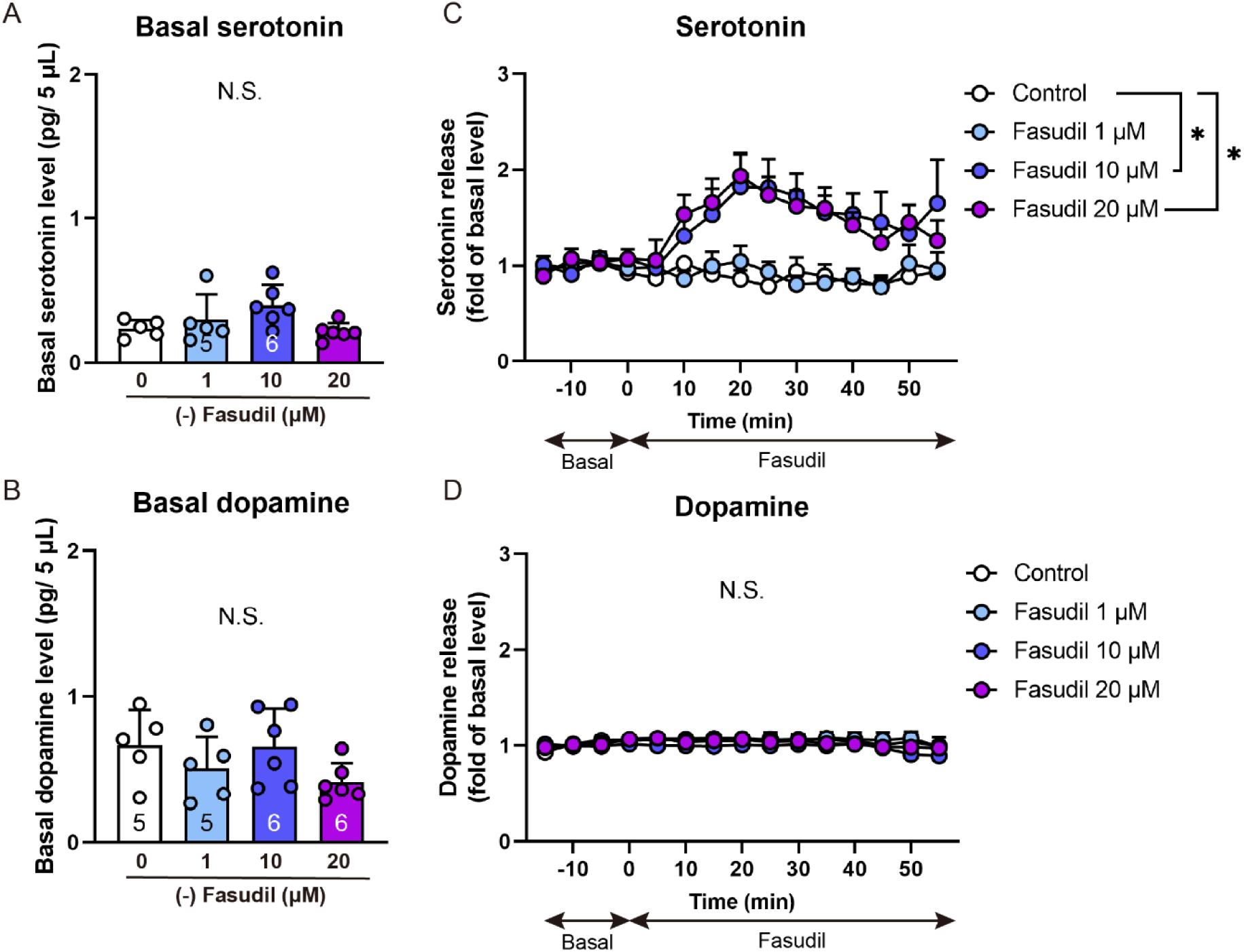
Fasudil perfusion into the NAc increased extracellular serotonin levels but not dopamine levels in mice A: Basal serotonin release before drug treatment. B: Basal dopamine release before drug treatment. C: Serotonin release during fasudil perfusion into the NAc of mice. D: Dopamine release during fasudil perfusion into the NAc of mice. Following the collection of the three baseline fractions (A, B), mice were perfused fasudil (1, 10, and 20 μM) into the NAc through the dialysis probe. Data represent the mean ± SEM (n = 5–6). *p < 0.05, significantly different from control (two-way repeated measures ANOVA). NAc, Nucleus accumbens; ANOVA, one-way analysis of variance

**Figure 3.**
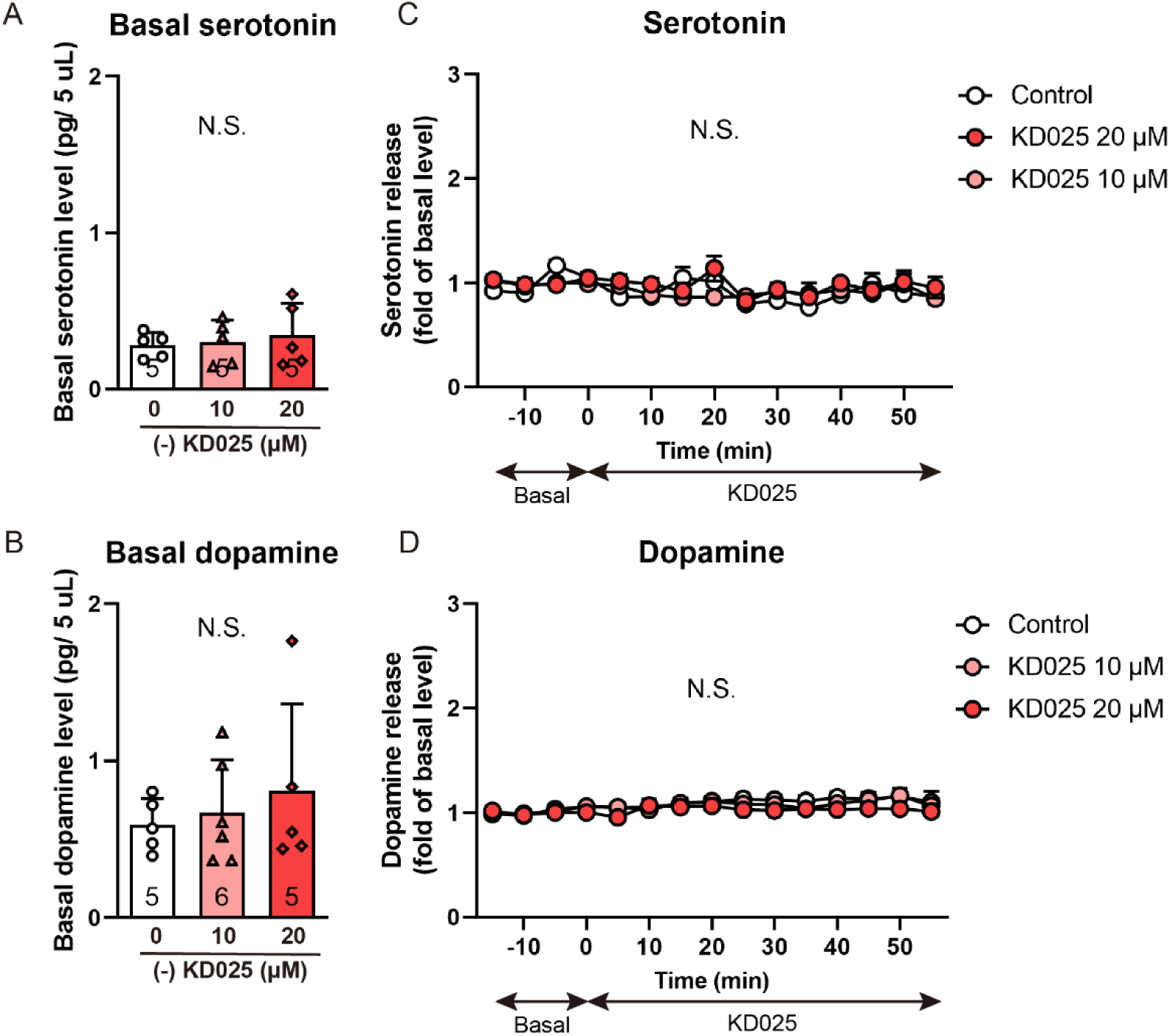
KD025 perfusion into the NAc had no effect on basal extracellular serotonin or dopamine levels in mice A: Basal serotonin release before drug treatment. B: Basal dopamine release before drug treatment. C: Serotonin release during KD025 perfusion into the NAc of mice. D: Dopamine release during KD025 perfusion into the NAc of mice. Following the collection of the three baseline fractions (A, B), mice were perfused KD025 (10 or 20 μM) into the NAc through the dialysis probe. Data represent the mean ± SEM (n = 5–6). NAc, Nucleus accumbens

**Figure 4.**
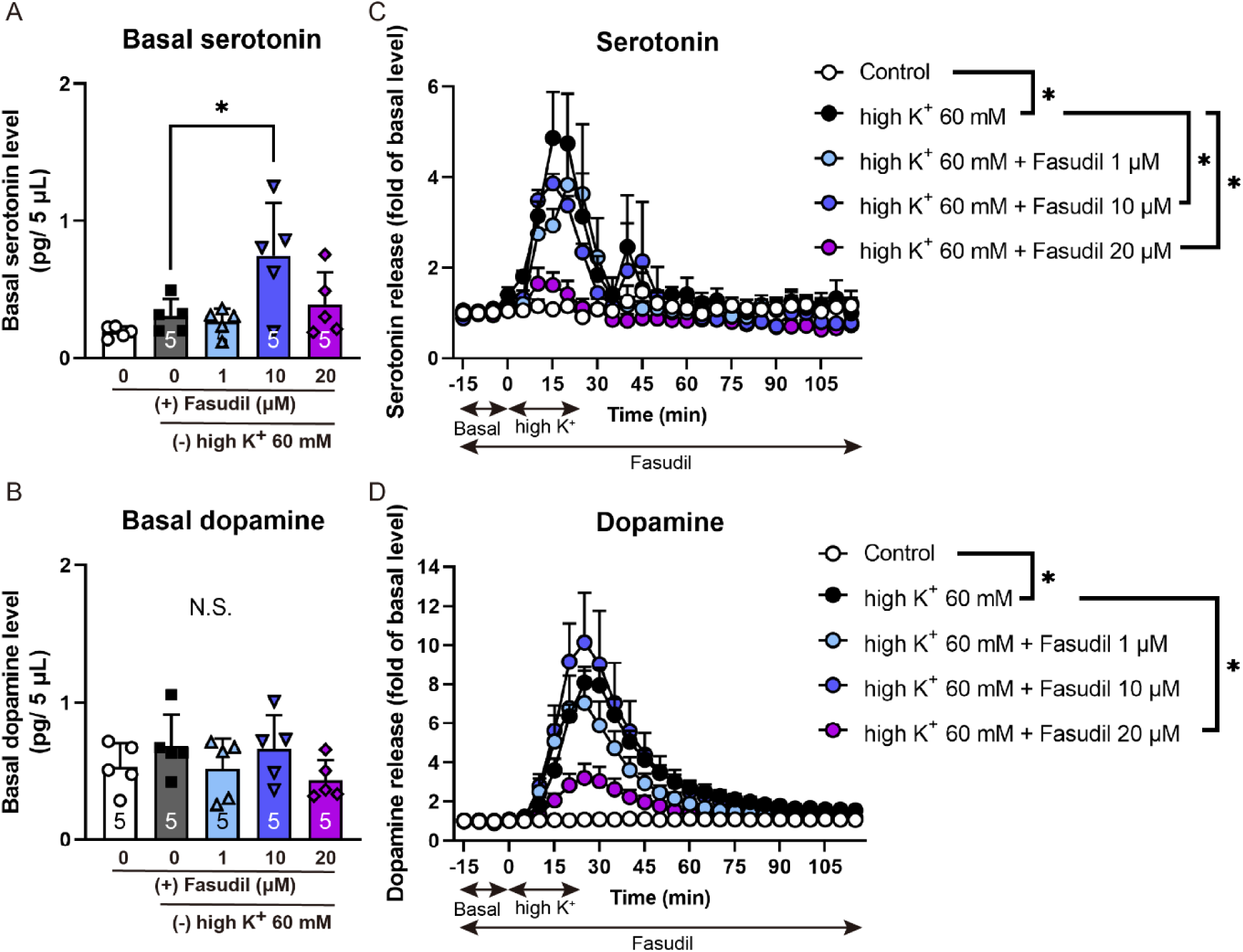
Fasudil perfusion into the NAc suppressed depolarization-induced serotonin and dopamine release in mice A: Basal serotonin release before stimulation. B: Basal dopamine release before stimulation. C: Depolarization-induced serotonin release in the NAc during fasudil perfusion in mice. D: Depolarization-induced dopamine release in the NAc during fasudil perfusion in mice. Following the collection of the three baseline fractions (A, B), high-K^+^-containing aCSF (60 mM KCl) with fasudil (1, 10, and 20 μM) was perfused into the NAc for 20 min through the dialysis probe. Data represent the mean ± SEM (n = 5). *p < 0.05, significantly different from control (two-way repeated measures ANOVA). NAc, Nucleus accumbens; ANOVA, one-way analysis of variance

### Histochemistry

Mice were perfused intracardially using ice-cold 0.1 M phosphate-buffer (PB) followed by 4% PFA in 0.1 M PB under anesthesia with isoflurane. The brains were postfixed with the same fixative and cryoprotected using 20% sucrose followed by 30% sucrose in 0.1 M PB. Frozen 30-μm sections were cut coronally using a cryostat (CM1860; Leica, Wetzlar, Germany). Cryosections were fixed with 4% paraformaldehyde in 0.1 M PB for 5 min and permeabilized with 0.3% Triton X-100/phosphate-buffered saline (PBS) for 10 min. Sections were incubated with Hoechst 33342 (346–07951, 1:2000, Dojindo, Kumamoto, Japan) at 25 °C for 1 h. After washing in PBS, the sections were mounted on an adhesive silane (MAS)-coated glass slide (Matsunami, Osaka, Japan) with a Fluorescent Mounting Medium (Dako, Santa Clara, CA, USA) and coverslip. Fluorescence images were captured using a BZ-X800 bright-field microscope (KEYENCE, Hyogo, Japan) equipped with a 2× objective lens.

### Statistical analysis

All data are expressed as the mean ± SEM. Statistical analyses were performed using GraphPad Prism (version 9; RRID:SCR_002798; GraphPad Software, San Diego, CA, USA). Statistical significance (P < 0.05) was determined using one-way analysis of variance (ANOVA) and two-way repeated-measures ANOVA for multigroup comparisons. Tukey’s multiple comparison test was used for post hoc comparisons. The sample size for each experiment was determined based on previous studies [31]. Detailed information on the statistical analyses is presented in Table S1.

## Results

Fasudil perfusion into the NAc increased extracellular serotonin levels but not dopamine levels in mice

First, we investigated the effect of a nonselective Rho-kinase inhibitor, fasudil, on the basal extracellular levels of serotonin and dopamine in the NAc. We confirmed that the basal serotonin and dopamine levels in the NAc were not different among the groups before fasudil treatment (Fig. 2A and B). Fasudil (10 and 20 μM) perfusion significantly increased the extracellular serotonin levels in the NAc 20 min post-treatment (Fig. 2C), but not dopamine levels (Fig. 2D).

KD025 perfusion into the NAc had no effect on basal extracellular serotonin or dopamine levels in mice

Next, we investigated the effects of a selective Rho-kinase 2 inhibitor, KD025, on basal extracellular serotonin and dopamine levels in the NAc. The basal extracellular serotonin and dopamine levels in the NAc were comparable among the groups before KD025 treatment (Fig. 3A and B). In contrast to the effect of fasudil (Fig. 2), KD025 perfusion into the NAc did not affect the extracellular serotonin or dopamine levels (Fig. 3C and D). These results suggested the involvement of Rho-kinase 1, but not Rho-kinase 2, in the fasudil-induced increase of the basal extracellular serotonin levels in the NAc.

Fasudil perfusion into the NAc suppressed depolarization-induced serotonin and dopamine release in mice

To clarify the role of Rho-kinase in depolarization-induced neurotransmitter release in the NAc, we investigated the effect of fasudil on high-K^+^-induced serotonin and dopamine release in the NAc of mice. Because fasudil itself increased the basal extracellular levels of serotonin (Fig. 2C), the basal levels of serotonin during pretreatment with fasudil (10 μM), but not of dopamine, were significantly higher than those following pretreatment with the vehicle (Fig. 4A and B). Perfusion of high-K^+^-containing aCSF into the NAc for 20 min significantly increased both serotonin and dopamine release compared with those in the control group (Fig. 4C and D). Notably, fasudil (10 and 20 μM) significantly suppressed both the high-K^+^ (60 mM)- induced serotonin (Fig. 4C) and dopamine release (Fig. 4D) in the NAc in a dose-dependent manner. These results suggested that Rho kinases play a pivotal role in depolarization-induced serotonin and dopamine release in the NAc.

KD025 perfusion into the NAc suppressed depolarization-induced serotonin release but not dopamine release in mice

Finally, we investigated whether KD025 perfusion suppresses depolarization-induced serotonin and dopamine release in the NAc. The basal serotonin and dopamine levels in the NAc were comparable between the groups (Fig. 5A and B). Notably, KD025 perfusion (10 and 20 μM) selectively suppressed the high-K^+^ (60 mM)-induced serotonin, but not dopamine, release in the NAc in a dose-dependent manner (Fig. 5C and D). These results suggested that Rho-kinase 2 plays a pivotal role in the depolarization-induced serotonin release in the NAc.

**Figure 5.**
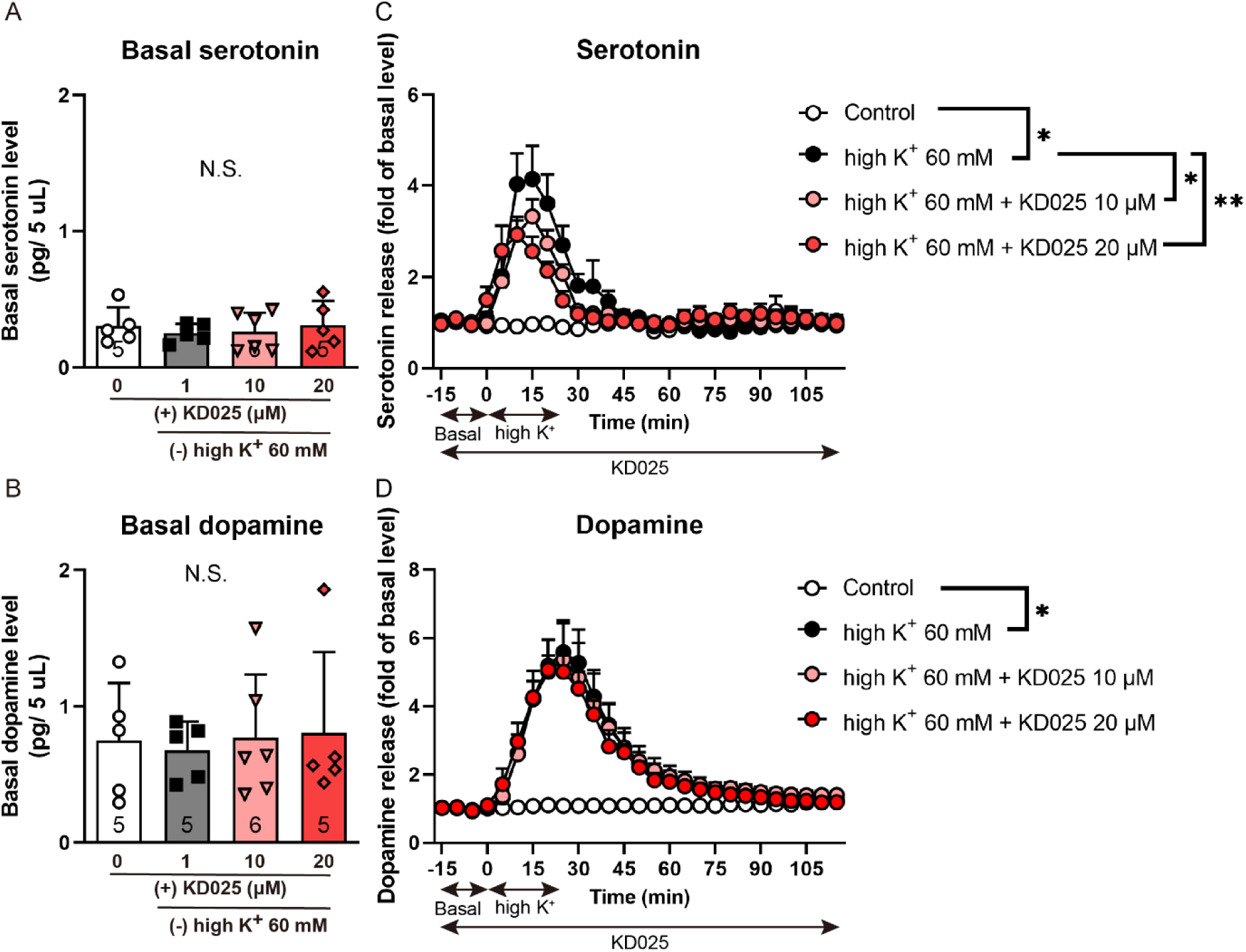
KD025 perfusion into the NAc selectively suppressed depolarization-induced serotonin release but not dopamine release in mice A: Basal serotonin release before stimulation. B: Basal dopamine release before stimulation. C: Depolarization-induced serotonin release in the NAc during KD025 perfusion in mice. D: Depolarization-induced dopamine release in the NAc during KD025 perfusion in mice. Following the collection of the three baseline fractions (A, B), high-K^+^-containing aCSF (60 mM KCl) with KD025 (10 or 20 μM) was perfused into the NAc for 20 min through the dialysis probe. Data represent the mean ± SEM (n = 5–6). *p < 0.05, **p < 0.01, significantly different from control (two-way repeated measures ANOVA) NAc, Nucleus accumbens; ANOVA, analysis of variance

## Discussion

Rho-kinase is a serine/threonine kinase that phosphorylates various substrates involved in regulating actin filament assembly [14]. Actin is involved in the translocation of synaptic vesicles to a readily releasable pool, exocytosis, and endocytosis [32–34].

Herein, we demonstrated that fasudil perfusion into the NAc increases basal extracellular serotonin levels (Fig. 2). Hence, Rho-kinase may play a role in the regulation of basal extracellular serotonin levels in the brain, although some studies have reported the effects of Rho-kinase on extracellular neurotransmitter levels [15,22]. In rat brain synaptosomes, the nonselective Rho-kinase inhibitor Y-27632 increases glutamate release without depolarizing the plasma membrane and induces a Ca^2+^-independent increase in acridine orange fluorescence, a pH-sensitive dye, suggesting an increase in the vesicular release of neurotransmitters [22]. Notably, acridine orange fluorescence increased throughout the assay, as shown in Fig. 2C, without a decline in fluorescence observed with KCl, as shown in Figs. 4C and 5C [22]. As both Y-27632 and fasudil increase the release of different neurotransmitters, such as glutamate and serotonin, these data suggest that Rho-kinase inhibitors decrease the pH gradient in synaptic vesicles and lead to the leakage of neurotransmitters to the cytosol and their pumping out by plasma membrane transporters [22]. In addition, Rho-kinases mediate the docking and fusion of vesicles and synaptic vesicle endocytosis [15,20,35]. These findings suggest that fasudil regulates basal extracellular serotonin levels via a Ca^2+^-independent mechanism without causing depolarization.

Rho-kinases have been reported to regulate dopamine uptake via mediation of dopamine transporter expression on the cell surface [36–39]. The present study demonstrated that fasudil selectively increases the basal extracellular serotonin levels in the NAc (Fig. 2C and D). Therefore, Rho kinases may mediate the localization of dopamine and serotonin transporters in different ways. However, the roles of Rho-kinases in the localization of dopamine transporters are controversial [36–39] and no reports are available on the relationship between Rho-kinases and serotonin transporter localization in neurons. To clarify the mechanism by which fasudil increases the basal extracellular serotonin levels in the NAc, we should further investigate the localization of dopamine and serotonin transporters in our model.

The Ca^2+^-dependent protein kinase CaMK2 phosphorylates ARHGEF2 to enhance its RhoGEF activity, leading to the activation of RhoA/Rho-kinase [40]. Indeed, high-K^+^ stimulation increases the phosphorylation of myosin phosphatase target subunit-1, a substrate of Rho-kinase [41]. We demonstrated that the perfusion of fasudil, a nonselective Rho-kinase 1 and 2 inhibitor, and KD025, a selective Rho-kinase 2 inhibitor, into the NAc suppressed depolarization-induced serotonin levels (Figs. 3 and 4). This strongly suggested that Rho-kinase 2 plays a pivotal role in depolarization-induced serotonin release in the NAc in vivo. However, reports on the effects of Rho-kinase on depolarization-induced neurotransmitter release in vitro and ex vivo are controversial [17–21,23,24]. Y-27632 suppresses high-K^+^-induced norepinephrine synthesis and release in PC12 cells [18] and electrical field stimulation (EFS)- induced acetylcholine release [17]. In contrast, Y-27632 increases the high-K^+^-induced exocytosis of FM 2-10 in rat cortical synaptosomes and EFS-induced outflow of [^3^H]-choline from airway cholinergic nerves [19,23]. Because Rho-kinase has various substrates and downstream molecules associated with synaptic function [21,23], it likely plays multiple roles in Ca^2+^-dependent exocytosis depending on the conditions.

However, overexpression of the constitutively active mutant of Gα_12_ or Gα_13,_ which induced RhoA/Rho-kinase activation, inhibited the high-K^+^-induced dopamine release in PC cells, but did not affect the high-K^+^-induced increase in intracellular Ca^2+^ concentration [24]. As Rho-kinase is downstream of CaMK2, it may mediate the downstream signaling of the depolarization-induced increase in Ca^2+^ concentration. To clarify the mechanism by which Rho-kinase mediates Ca^2+^-dependent exocytosis, further evaluation of the Ca^2+^-CaMK2-Rho-kinase signaling is required.

Our data showed the different effects of fasudil and KD025 on dopamine and serotonin release. In particular, Rho-kinase 1 may be associated with depolarization-induced dopamine release, whereas Rho-kinase 2 may be associated with depolarization-induced serotonin release. These results suggested that Rho-kinases 1 and 2 may play different roles in neurotransmitter release from serotonin and dopamine neurons projecting to the NAc. Fasudil and KD025 showed similar effects on spine density in the mPFC and cognitive dysfunction in schizophrenia mouse models but showed distinct effects on the impairment of pre-pulse inhibition [10,12]. In addition, Rho-kinase 1 or 2 knockout mice showed comparable spine density, whereas apical spine length was decreased in Rho-kinase 1 knockout mice but increased in Rho-kinase 2 knockout mice [28]. Notably, these knockout mice showed a distinct effect on the phosphorylation levels of the myosin light chain, a Rho-kinase substrate, in synaptosomes [28]. These suggested that Rho-kinases 1 and 2 have distinct functions in neurons.

Rho-kinases 1 and 2 are 64% homologous, with their kinase domains exhibiting the highest homology (92%) and similar kinase activity [25–27]. However, the expression of these subtypes varies among tissues [27]. Rho-kinase 1 is predominantly expressed in the blood cells and thymus, whereas Rho-kinase 2 is abundantly expressed in the brain and heart of mice and humans [25,42]. In the brain, Rho-kinases are expressed in neurons at both pre- and postsynaptic terminals [12,23,43,44]. As their expression patterns in neurons are distinct at the neonatal stage [20], the localization of Rho kinases 1 and 2 in neurons may also differ at the adult stage. In addition, their distribution patterns in presynaptic dopamine and serotonin neurons remain unclear. To clarify the functional differences between Rho-kinases 1 and 2 in neurons, evaluating the presynaptic localization of Rho-kinases 1 and 2 in each neuronal subtype is necessary.

The present study had two limitations. First, currently no selective Rho-kinase 1 inhibitors are available; thus, we could not clarify the selective role of Rho-kinase 1 in neurotransmitter release. Second, although fasudil is widely used as a Rho-kinase inhibitor (IC_50_ of Rho-kinase = 10.36 μM; IC_50_ of Rho-kinase 2 = 0.158 μM), it inhibits other kinases such as PKA (IC_50_ 4.58 μM), PKG (IC_50_ 1.65 μM), and CaMK2 (IC_50_ 6.70 μM) [45,46]. Likewise, KD025 selectively inhibits Rho-kinase 2 (IC_50_ of Rho-kinase >10 μM; IC_50_ of Rho-kinase 2 = 0.059 μM) and casein kinase 2 (IC50 = 50 nM) [47,48]. Therefore, the present study cannot exclude other candidate molecules that may be involved in the effects of fasudil and KD025. To obtain solid evidence of the physiological roles of Rho-kinases 1 and 2 in neurotransmitter release, studies using genetic methods, such as using conditional Rho-kinase knockout mice, are required.

## Conclusion

In conclusion, our results suggest the following: Rho-kinases 1 and 2 play pivotal roles in the regulation of neurotransmitter release in the NAc, Rho-kinase 1 may be associated with depolarization-induced dopamine release, and Rho-kinase 2 may be specifically associated with depolarization-induced serotonin release.

## Acknowledgments

Fasudil hydrochloride was kindly provided by Asahi Kasei Pharma Corp. (Tokyo, Japan). The authors wish to acknowledge the Division for Medical Research Engineering, Nagoya University Graduate School of Medicine, for the use of the BZ-X800 Bright-Field Microscope (KEYENCE).

## Conflicts of Interest

This study was partially funded by Sumitomo Pharma. Fasudil hydrochloride was provided by Asahi Kasei Pharma Corp..

## Funding Information

This study was supported by the Japan Agency for Medical Research and Development (AMED) (JP21wm0425007) and the Japan Society for the Promotion of Science (JSPS) KAKENHI (JP23H02669, JP23K19425, and JP24K18365).

## Data Availability Statement

The data that supports the findings of this study are available in the supplementary material of this article.

## Approval of the research protocol by an Institutional Reviewer Board

n/a

## Informed Consent

n/a

## Registry and the Registration No. of the study/trial

n/a

## Animal Studies

All procedures were conducted in accordance with the guidelines established by the Institutional Animal Care and Use Committee of Nagoya University, Guiding Principles for the Care and Use of Laboratory Animals approved by the Japanese Pharmacological Society, and Guide for the Care and Use of Laboratory Animals by the National Institutes of Health of the United States.

## Author Contributions

**Rinako Tanaka:** Writing – original draft, Investigation, Methodology, Data curation, Conceptualization, Funding acquisition. **Taku Nagai:** Conceptualization, Writing – review and editing**. Toshitaka Nabeshima:** Conceptualization, Writing – review and editing**. Kozo Kaibuchi:** Conceptualization, Writing – review and editing, **Norio Ozaki:** Funding acquisition, Writing – review and editing. **Hiroaki Ikesue:** Writing – review and editing. **Hiroyuki Mizoguchi:** Supervision, Writing – review and editing. **Kiyofumi Yamada:** Conceptualization, Supervision, Writing – review and editing, Funding acquisition.

## Notes

**Funding statement** This study was supported by the Japan Agency for Medical Research and Development (AMED) (JP21wm0425007) and the Japan Society for the Promotion of Science (JSPS) KAKENHI (JP23H02669, JP23K19425, and JP24K18365).

**Conflict of interest disclosure** This study was partially funded by Sumitomo Pharma. Fasudil hydrochloride was provided by Asahi Kasei Pharma Corp.

